# Esearch3D: Propagating gene expression in chromatin networks to illuminate active enhancers

**DOI:** 10.1101/2022.08.04.502774

**Authors:** Maninder Heer, Luca Giudice, Claudia Mengoni, Rosalba Giugno, Daniel Rico

## Abstract

Most cell type-specific genes are regulated by the interaction of enhancers with their promoters. The identification of enhancers is not trivial as enhancers are diverse in their characteristics and dynamic in their interaction partners. Currently, enhancer-associated features such as histone modifications, co-activators or bi-directional transcription are used in lieu of any definitive and universal enhancer feature. We present Esearch3D, a new approach that leverages network theory approaches to identify active enhancers. Our work is based on the fact that enhancers act as a source of regulatory information to increase the rate of transcription of their target genes and that the flow of this information is mediated by the folding of chromatin in the three-dimensional (3D) nuclear space between the enhancer and the target gene promoter. Esearch3D reverse engineers this flow of information to calculate the likelihood of enhancer activity in intergenic regions by propagating the transcription levels of genes across 3D-genome networks. Regions predicted to have high enhancer activity are shown to be enriched in annotations indicative of enhancer activity. These include: enhancer-associated histone marks, bi-directional CAGE-seq, STARR-seq, P300 and RNA polymerase II ChIP-seq, and expression quantitative trait loci (eQTL). Esearch3D successfully leverages the relationship between chromatin architecture and global transcription and represents a novel approach to predict active enhancers and understand the complex underpinnings of regulatory networks. The method is available at: https://github.com/InfOmics/Esearch3D.

## Introduction

The non-coding genome contains a diverse collection of regulatory regions including non-coding RNAs, silencers, insulators and cis/trans regulatory elements. Transcriptional enhancers, a class of distal cis regulatory elements, are of particular interest as they are able to direct the cell-type specific transcriptional activity of genes (1). Defining enhancers and disentangling their spatio-temporal localisation with promoters is important in the context of understanding a wide variety of biological phenomena such as evolution, homeostasis and disease (2, 3). However, current definitions of enhancers are broad and non-specific, while their mode of action is poorly understood. Despite the lower costs and advances in whole genome sequencing, problems persist in attempts to consolidate the link between enhancers, non-coding genetic variation and protein coding genes for a great number of diseases. The enhancer characteristics underpinning these difficulties can be broadly summarised by the following points:

1. *Enhancers are promiscuous*. A single enhancer can regulate multiple genes (4).
2. *Enhancers can be redundant*. Perturbing one enhancer can have a minimal effect on the expression of a gene that may rely on several enhancers (5–7).
3. *Enhancers interactions are complex and dynamic*. Enhancers can be located up to, and in some cases beyond, one megabase away from their cognate gene (8) and interact with gene promoters in specific three-dimensional spatio-temporal patterns (9).
4. *Enhancer sequence composition is highly heterogeneous*. There is no known conserved enhancer sequence that universally defines all enhancers (2).

To address these difficulties, current methods to find enhancers have sought to identify them through their associated features. One of the most common features are histone modifications. H3K4me1 is associated with enhancers in the active (when coexisting with H3K27ac), inactive and primed (when coexisting with H3K27me3) states (10). Other marks such as H3K4me3 (11), H3K79me2 (12), as well as H3K64ac, H3K122ac and H4K16ac (13) have also been found to be enriched at histones proximal to distinct sets of enhancer sequences. Currently, these marks can be used to identify putative enhancers but they are correlative and not universally present in all experimentally validated enhancers (13).

Beyond histone marks, coactivators such as CREB-binding protein (CBP) and P300 have also been proven to be a prerequisite for the activity of some enhancers by their acetyltransferase activity at H3K27 and recruitment of RNA polymerase II (RNAPII) (14, 15). Other methods that are currently used to identify enhancers include enhancer RNAs (eRNAs) that exhibit by bi-directional transcription and are quantified by cap analysis of gene expression (CAGE-seq) protocols; FANTOM has catalogued putative enhancers based on this assay (16). In addition, self-transcribing active regulatory region sequencing (STARR-seq) has become a popular high-throughput method to experimentally assess the transcriptional enhancing properties of the DNA sequences in putative enhancers (17–19). Finally, expression quantitative trait loci (eQTL) can be used to associate non-coding SNPs in putative enhancer sequences with changes in gene expression (20, 21).

Enhancer associated features can be used individually or in tandem to identify enhancers, albeit with varying levels of success (22). A way of improving such predictions is to integrate more data to better inform the model. The organisation of chromatin into higher order hierarchical structures such as A/B compartments, topologically associated domains (TADs) and enhancer promoter loops has been associated with orchestrating cell-type specific gene regulation by localising regulatory enhancers to promoters in specific spatio-temporal patterns (3, 23). These patterns of connectivity can be uncovered by methods known as chromosome conformation capture (3C) such as HiC, capture-HiC, ChIA-PET or HiChIP (24). These specific patterns of 3D chromatin conformation add yet another layer of complexity to the genome and represent a feature from which the spatio-temporal localisation of enhancers with promoters can be determined. Indeed, there are some pioneering predictive algorithms that have sought to leverage this information to link enhancers with their target genes in specific cell lines using 3C data, accessible chromatin information and active histone marks (25) or using deep learning models trained with massive collections of chromatin and gene expression data to predict distal regulatory regions from DNA sequence (26).

A caveat of using 3C data from bulk cell populations is that it provides a snapshot of the most frequent chromatin interactions and is therefore insensitive to the dynamic and transient nature of many enhancer-promoter interactions. Additionally, single cell HiC approaches are still only capturing a very small proportion of the interactions (27) and do not allow the comprehensive detection of promoter-enhancer interactions at adequate resolutions. There is increasing evidence that well established enhancer-promoter contacts are not a prerequisite for all enhancer activity. For example transcriptional bursting has been demonstrated to enable the periodic transcription of genes (28) but the temporal dynamics of promoter-enhancer interactions appear to be uncoupled from those of transcription (29). Moreover, enhancer activity has also been observed with a distinct lack of enhancer-promoter proximity (30). These types of interactions are susceptible to be missed as, typically, 3C data is interrogated in a pairwise manner. Although microscopy approaches can address this they are limited by their low-throughput. Therefore, a useful way to capture these lost pairwise interactions is to represent 3C data as networks. In a network an interaction between loci A and C can be inferred through their common interaction with loci B that would otherwise be lost using established pairwise methods.

Representing these 3C data as networks has the potential to capture more chromatin interactions as well as uncover hidden trends beyond direct interactions such as cooperative regulatory subnetworks (**Figure 1**). Such advantages of network analysis have already been demonstrated. For example, Sandhu et al constructed a chromatin interaction network from RNAPII mediated interactions describing large scale organisation into chromatin communities with specific function (31). We were able to use chromatin interaction networks and assortativity measures to demonstrate how specific features of the chromatin are enriched in promoter-promoter contacts vs promoter-non-promoter contacts (32). Thibodeau and colleagues have demonstrated that H3K4me3 broad domains and super enhancers in chromatin interaction networks maintained unique and distinct topological properties that could be used to distinguish broad domains from promoters and super-enhancers from normal enhancers (33). Finally, recent reports have shown the usefulness of graph-based approaches to detect functional modules in 3C data (34–37).

**Figure 1.**
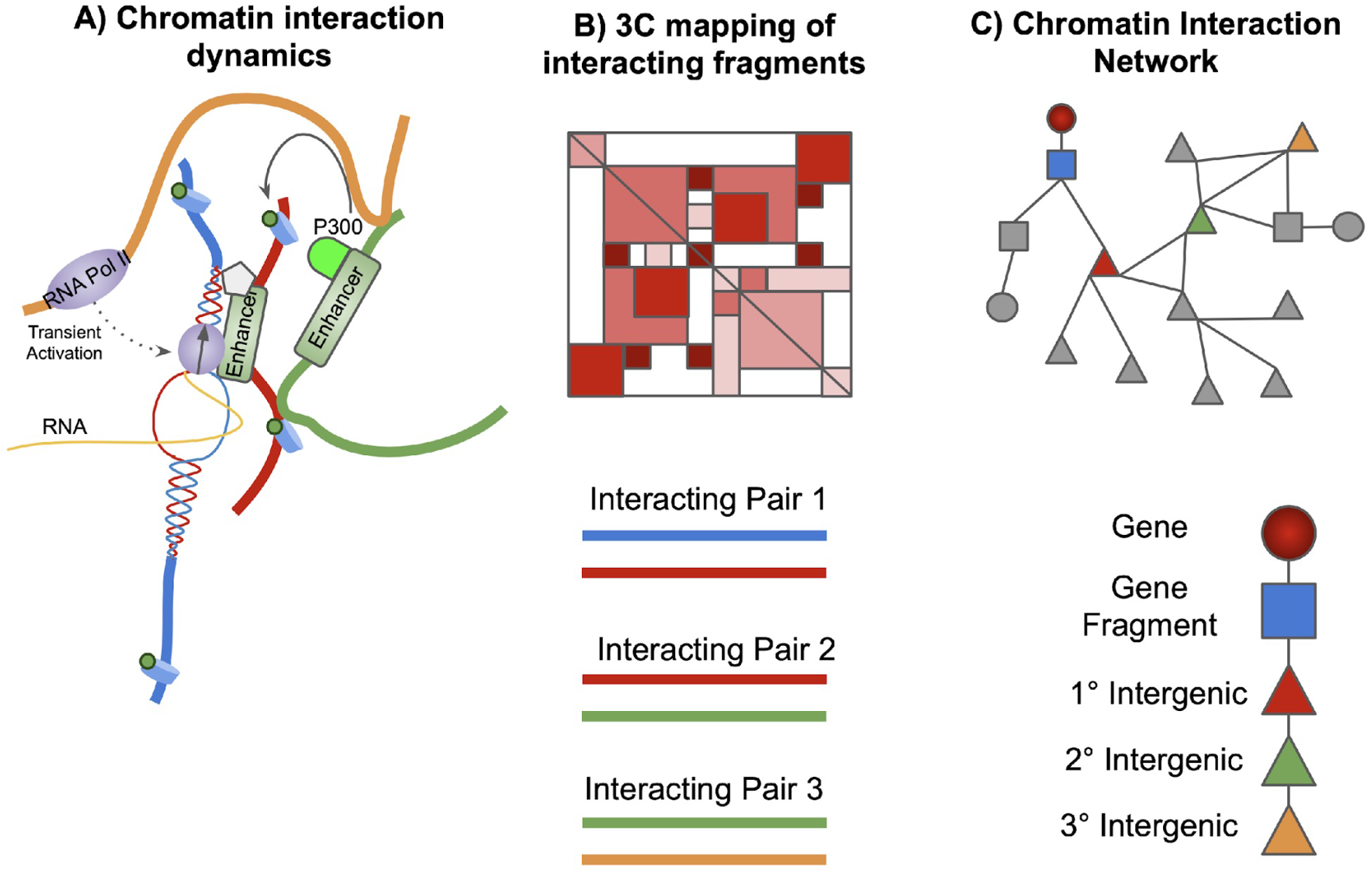
Schematic of converting chromatin structure datasets into networks. (**A)** Enhancers can influence the regulation of promoters directly and indirectly. (**B**) HiC and HiC-derived methods can capture chromatin interactions but only in a pairwise manner. (**C**) Representation of chromatin fragments as nodes in a network, with their interactions as edges, preserves indirect associations between interacting chromatin fragments and allows global system analyses.

It is becoming increasingly apparent that the global organisation of chromatin and the accumulation of perturbations across the genome can have subtle but tangible effects on gene expression of most expressed genes. An *omni-genic* model has been proposed, where the activity of almost any gene would affect that of almost every other one, as a result of the small world properties of multiple layers of regulatory networks (38). This model was proposed as an attempt to explain the multi-loci nature of many complex diseases such as Alzehimer’s (39) or Crohn’s disease (38) that has been uncovered by genome wide association studies (GWAS). Most expressed genes in a given cell type would be only a few steps from the nearest core disease gene and thus may have non-zero effects on disease. As these mechanisms likely act to influence the localisation of regulatory enhancers to gene promoters, a *pan-genomic* model based purely on 3D genome interactions has been later proposed to reconcile the strong effects of over-expressing transcription factors in changing cellular phenotypes with the data from eQTL studies (where genetic effects are small but widespread) (40). Focusing solely on the 3D genome network and transcription, this work shows how just changing the activity of one gene can dramatically change the topology of the networks and the overall transcriptional programme, showing how we cannot understand the regulation of a particular gene in isolation. However, none of the current methods to identify the enhancers regulating a particular gene take into account the expression of the other genes being transcribed.

Here, we describe our Esearch3D approach that exploits the global relationship between 3D genome and transcription to identify active enhancers in 3C data. Biologically, signals pertaining to transcriptional activation are transferred across the regulatory network from enhancers found in intergenic loci to genes in the form of transcription factors, cofactors and various transcriptional machinery such as RNAPII. Conceptually it can be described as a flow of information between enhancers and genes, mediated by the chromatin communication network. How and where this information is transmitted to and from is central to decoding the regulatory landscape of any gene and therefore, identifying enhancers. We show that by reverse engineering this flow of information we can identify intergenic regulatory enhancers using solely gene expression and 3D genomic data.

## Results

### Overview and conceptual description of Esearch3D

We have developed an R package called Esearch3D. Primarily, the tool relies on a bespoke unsupervised algorithm based on random walks to predict active enhancers using solely gene expression and chromatin interaction data. Enhancers are non-randomly located and connected in the network such that they have the potential to regulate cell type-specific gene expression. Esearch3D exploits this phenomenon to identify intergenic regions with regulatory functions. To achieve this, Esearch3D propagates the gene expression values of gene-containing nodes to all other nodes in the 3C-derived networks (DNaseI capture Hi-C and Promoter capture Hi-C) (see example with a toy network in **Figure 2**). The result is an imputed activity score (IAS) in intergenic nodes. Much like gene expression reflects the transcriptional activity of a gene, IAS reflects the enhancer activity of non-coding regions. This score is based on the propagation of all expressed genes throughout the whole network. It is therefore a function of the intergenic nodes distance (i.e. the number of steps) to all other genic nodes in the network, the topology of the network and the expression levels of all genes within the network. In addition to predicting enhancers, our method highlights the integral role of global chromatin organisation in mediating gene expression.

**Figure 2.**
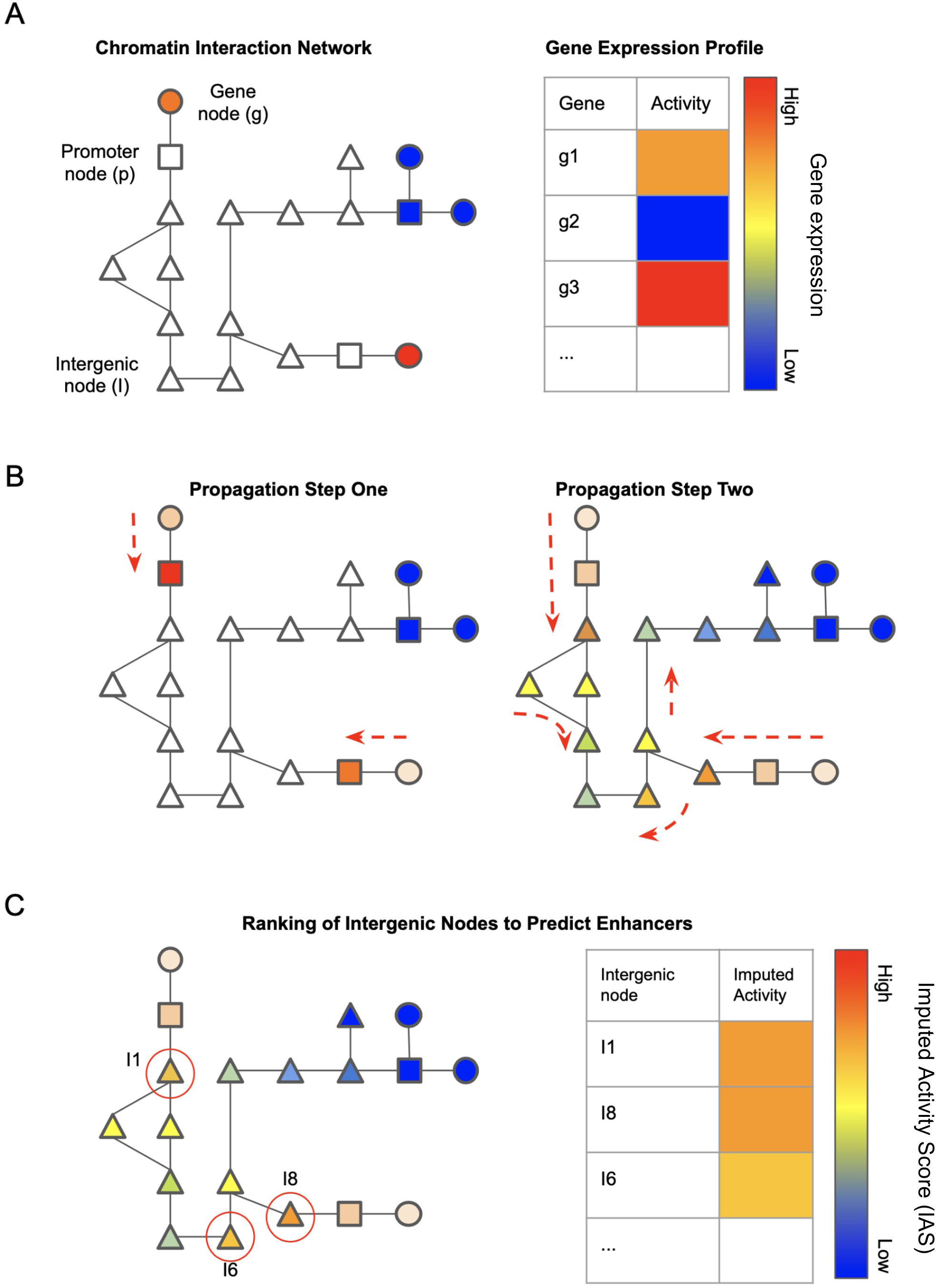
Schematic diagram of the network propagation used by Esearch3D to impute activity values at intergenic nodes. (**A**) Genes are mapped to nodes representing genic chromatin fragments. Each gene has an associated gene activity value determined by RNA-seq data. (**B**) Gene activity is propagated from gene nodes to genic chromatin nodes in propagation step one. Activity scores are then imputed in intergenic chromatin nodes by propagating the scores from genic chromatin nodes. (**C**) Ranking of intergenic nodes by the imputed activity score to identify high confidence enhancer nodes.

It is important to note that the integration of gene expression data and nodes in 3C-derived networks is not trivial, due to the discrepancies in 3C fragment resolutions - the genomic fragments can contain more than one gene and genes can expand into multiple fragments. Our method addresses this by incorporating genes as distinct additional nodes within the chromatin interaction network where they are connected by an edge to the node(s) that the body of their respective genes map to (**Figure 2**). Each gene node also contains an associated RNA-seq value that reflects the transcriptional activity (expression) of the gene. The expression profile of the genes serves as the *a priori* information that is to be propagated across the network (**Figure 2A**). Our approach uses the function F defined by Zhou et al. (41) to propagate gene expression across the chromatin interaction network (see **Material and Methods** and **Supplementary Figure 1**). To provide some intuition as to how this method works consider the following example. At a genic node a walker starts with some amount of information. In this case the value given by the gene expression. The walker then has equal probability to transition to any of its direct neighbours and carry a proportion of the gene expression value to that node. The amount of gene expression that can transition is determined by the insulating parameter alpha α. For example, where α = 0.8 only 20% of the original gene expression value can transition. Low values of α were chosen as genes are unlikely to be highly regulated by nodes very far away. The other parameter corresponds to the number of iterations and influences how many times this process takes place. A low number of iterations limits the amount of gene expression that can be carried to neighbouring nodes which can lead to these nodes obtaining similar values making it more difficult to distinguish their relative importance to the gene node of interest.

The expression profiles are propagated from the gene nodes into the genic nodes in propagation step one (**Figure 2B**). This facilitates the unbiased assignment of gene expression profiles to genes that map to multiple nodes. The gene expression profiles are then propagated to the rest of the 3D genome network in propagation step two (**Figure 2B**). Following propagation, the intergenic nodes (genomic regions not overlapping with genes) receive an imputed activity score (IAS) reflecting the likelihood of enhancer activity based on the distance to all genes in the network and the expression levels of these genes (**Figure 2C).**

### Imputed activity scores at intergenic nodes are higher in active enhancer nodes

Esearch3D infers the likelihood of enhancer activity at an intergenic locus with IAS following the propagation of gene expression. Esearch3D requires two inputs: 3C data which is modelled as a network and gene expression data. We generated our first set of results using the high-resolution DNaseI Capture Hi-C generated by Joshi and co-workers (42) in mouse embryonic stem cells (mESCs). This dataset was chosen owing to a diverse set of publicly available canonical enhancer associated features in mESCs, which can be used to validate our predictions (**Figure 3A**). DNaseI capture HiC targets regions of the mouse genome that are hypersensitive to DNaseI (DHS regions), enriching for chromatin accessible regions. From this DHS Capture HiC (DHS-CHiC) data, we created a chromatin interaction network, where each genomic fragment from this data set is represented as a node and each fragment-fragment interaction as an edge (**Figure 1**). The DHS-CHiC network consists of a total of 162,615 nodes connected by 787,879 edges with an average degree of 9.69. The chromatin interaction network is split across 776 connected components, however the majority of the nodes (97.7%) are members of the major connected component. We then identified genic nodes by mapping Ensembl-defined genes to the nodes (see **Material and Methods** and **Supplementary Table 1-3**). We found that 36% of the nodes in the chromatin interaction network were labelled as intergenic; that is to say that there are no overlaps of the gene body (including UTRs, exons and introns). Although introns also have many enhancers (see (43), for example), we focused here in the prediction of enhancers not overlapping with genes.

**Figure 3.**
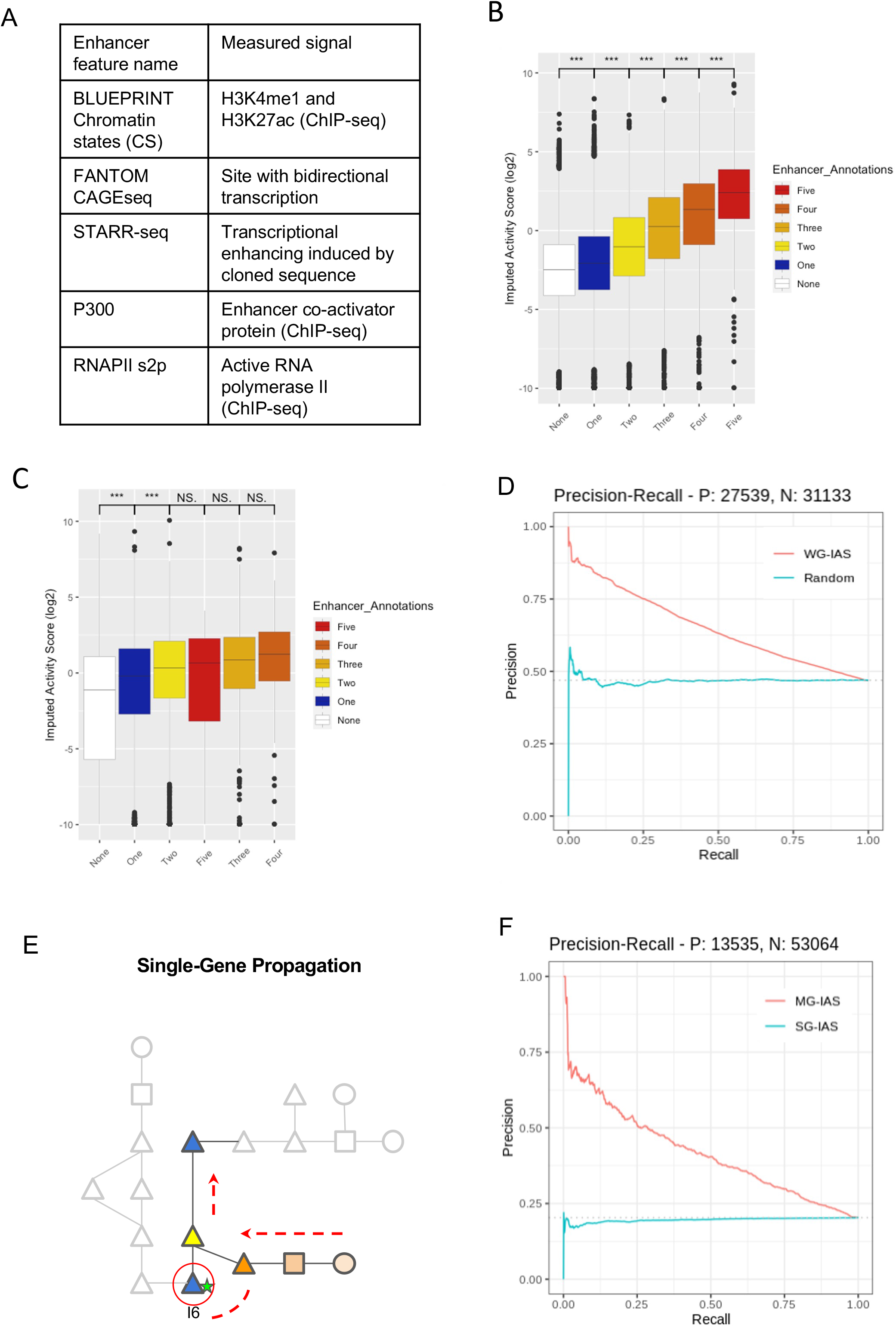
Enrichment of Esearch3D imputed activity score in enhancer annotated nodes. (**A**) Enhancer associated features in mouse embryonic stem cells (see Material and Methods for details). (**B**) Intergenic nodes in the mESC DHS-PCHiC network are stratified into enhancer and non-enhancer groups based on the number of enhancer-associated features that were annotated previously. The imputed activity score (IAS) is then plotted along the X-axis for each group of nodes. Boxplots are ordered by the median of each group and asterisks indicate significance in Wilcoxon tests (* = p.value < 0.05, ** = p.value < 0.005 *** = p.value < 2.2e-16). (**C**) IAS in the mESC P-CHiC network by number of enhancer annotations. (**D**) Precision recall curves measuring the overall performance of IAS in the prediction of enhancer nodes containing a minimum of 1 enhancer feature in the real (red line) and randomised (blue) networks. **(E)** A schematic of how single-gene propagation differs in the relative imputed activity scores. (**F**) Precision recall curves for the multi-gene propagation (red) vs the single gene propagation (blue). For single gene propagation we iteratively propagated the top 100 genes by expression and tested the predictive performance of MG-IAS on the intergenic nodes that receive a score. The multi-gene propagation performs with an AUPRC of 0.41 while the single gene propagation AUPRC is 0.19 over a baseline of 0.20.

To validate the efficacy of IAS as a predictive feature of intergenic enhancers, nodes were then labelled using the following enhancer associated features: chromatin states derived from histone mark ChIP-seq experiments, P300, RNAPII s2p elongating variant (44), STARR-seq (45) and CAGE-seq data from the FANTOM project (16, 46) (see **Materials and Methods** and **Figure 3A**). In the mESC DHS-CHiC network we found that in nodes that contain at least one enhancer associated feature there was a significant difference in the median value of IAS in enhancer nodes compared to non-enhancer nodes (Wilcoxon test p-value < 2.2e-16). Non-genic nodes were then stratified into 6 groups based on the different combinations of the five enhancer labels as well as the absence of any labels. We found that the median value of IAS in each group increased as the number of enhancer labels increased. While the median IAS from nodes devoid of any enhancer annotations to the group labelled with all 5 annotations increased by a factor of 46 (**Figure 3B**). These results show that the value of IAS is higher in intergenic enhancer nodes compared to non-enhancer nodes. Furthermore, IAS is found to be higher in nodes that contain an increasing number of enhancer features (**Figure 3B**).

We then applied Esearch3D to an alternative 3D genome capture data type by extending the analysis to Promoter-Capture HiC (PCHi-C) derived interactions for mESCs. The resulting network showed that PCHi-C results in a higher number of connected components resulting in a more disconnected network than the DHS-CHiC network (**Supplementary Tables 1-2**). Despite this, results are consistent with our previous observations using the DHS-CHiC network with higher IAS in enhancer nodes and significant increases in the median value of IAS between each group (Wilcoxon rank sum test p.value = < 2.2e-16, **Figure 3C**). IAS shows a minimum 14-fold enrichment overall in the high confidence enhancer nodes compared to the non-enhancer nodes for the mESC PCHi-C network.

These results show that Esearch3D can be used to approximate enhancer activity in chromatin fragments derived from DHS-CHiC and PCHi-C experiments. This is achieved independently from any *a priori* knowledge of enhancer associated features, using only gene expression and 3C contact data and without the need for model parameterisation. These results show that Esearch3D can be used to leverage the intrinsic relationship between the chromatin topology and gene expression to identify which nodes harbour enhancers.

### Imputed activity scores can be used to classify enhancer nodes

Following the observations of IAS enrichment in enhancer nodes, we then used IAS to classify nodes as enhancers and non-enhancers. Using the canonical enhancer associated features (**Figure 3A**) as our ground truth we plotted the precision-recall curves for the DHS-CHiC network and determined the area under the precision-recall curve (AUPRC, **Material and Methods**). IAS is able to correctly classify an enhancer node containing at least one of the five enhancer associated features with an AUPRC of 0.650 over a baseline of 0.469 (**Figure 3D**). The baseline is calculated as the ratio of enhancer nodes (n = 27,539) divided by the total number of intergenic nodes (n = 58,672). This is equal to a performance of 0.181 over baseline. For the mESC PCHi-C network there is a 0.095 increase over the baseline (**Supplementary Figure 2**), a 0.086 decrease when compared to the mESC DHS-CHiC network. The differences observed between the two capture methods are likely due to the smaller representation of the mESC genome by PCHi-C compared to a DNaseI capture that covers 50% more of the total genome. This has two main effects. One is the number of potential enhancers that are captured; the proportion of intergenic nodes in the DHS-CHiC network labelled as enhancers is 50% compared to 43% for the PCHi-C network. The second is that a lower coverage capture of the genome by the PCHi-C results in the capture of fewer enhancers across the genome. In the PCHi-C network we observe a lower average degree of 2.61 compared to 9.69 for the DNaseI network (**Supplementary Table 2**). We note that this does not drastically alter the density of the network which calculates the ratio between the number of actual connections vs the number of potential connections; the number of potential connections is given by the binomial coefficient of the number of nodes. This suggests that the PCHi-C is not losing information about the topology of the network, rather it is sampling a smaller proportion of the real chromatin interaction network.

### Chromatin topology and gene expression are crucial features to identify enhancers

The ability of Esearch3D to predict enhancer nodes is based on the propagation of gene expression across the chromatin interaction network. This diffuses gene expression data from the genic nodes to the intergenic nodes. Our results suggest that enhancer nodes maintain distinct connectivity patterns with gene nodes compared to non-enhancer nodes such that they receive a higher value of IAS. Therefore, the quality of the predictions rely on both the gene expression data and the network topology.

To demonstrate this we tested how the performance of IAS as a predictive feature changes when the topology of the network is perturbed. To do so, we applied a degree preserving rewiring algorithm to shuffle the edges of the mESC DNaseI network to create an ensemble of 100 rewired networks. This maintains the number of connections a node has while shuffling the node it is connected to. We then ran Esearch3D to the ensemble of randomised networks to predict enhancer nodes. We then aggregated and compared the predictions with those of the original network. Results showed that following the randomisation the ability of the model to classify enhancer nodes was lost with an AUPRC = 0.49 over a baseline of 0.47 (**Figure 3D**). This indicates that the spatial proximity of intergenic nodes with genic nodes is important for driving these predictions in line with evidence suggesting the role of transcriptional factories is a key determinant of gene expression (47).

There is increasing evidence that gene expression is regulated at a global level as outlined by the pan-genomic and omnigenic models. These models suggest that aggregate subtle changes in the expression of genes can influence the expression of others. For Esearch3D we modelled this process by propagating the expression of all genes across the network. To test the effects of this strategy we propagated the expression of a single gene of interest in order to identify the enhancers that directly regulate its expression. This approach is also available in the Esearch3D software. In this scenario, the expression profile of a single gene of interest was propagated across the chromatin interaction network. We carried out this test on the top 100 most highly expressed genes (112 when including ties). Results showed that the propagation of 112 single genes across the network reduced the performance of the model compared to that of a random classifier AUPRC = 0.19 (**Figure 3F**). Together, these results indicate that the global properties of the chromatin network and the global effects of gene expression are important features to identify enhancers.

### Esearch3D can be used to study the regulatory function of regions with eQTL SNPs

One of the main lessons gained from GWAS is that most disease associated small nucleotide polymorphisms (SNPs) lie within non-coding regions that are enriched in enhancers (48). Expression quantitative loci (eQTL) studies have shown how genetic variation across the genome can result in significant and quantifiable changes in gene expression (49). eQTL SNPs are defined by classifying individuals by that SNP genotype and calculating correlations with gene expression levels of genes in their linear genome proximity (**Figure 4A**). These approaches help to identify the cell types where disease-associated SNPs show an effect and to statistically associate non-coding SNPs with the potential gene targets, thereby aiding in the functional interpretation of GWAS hits outside genes (21).

**Figure 4.**
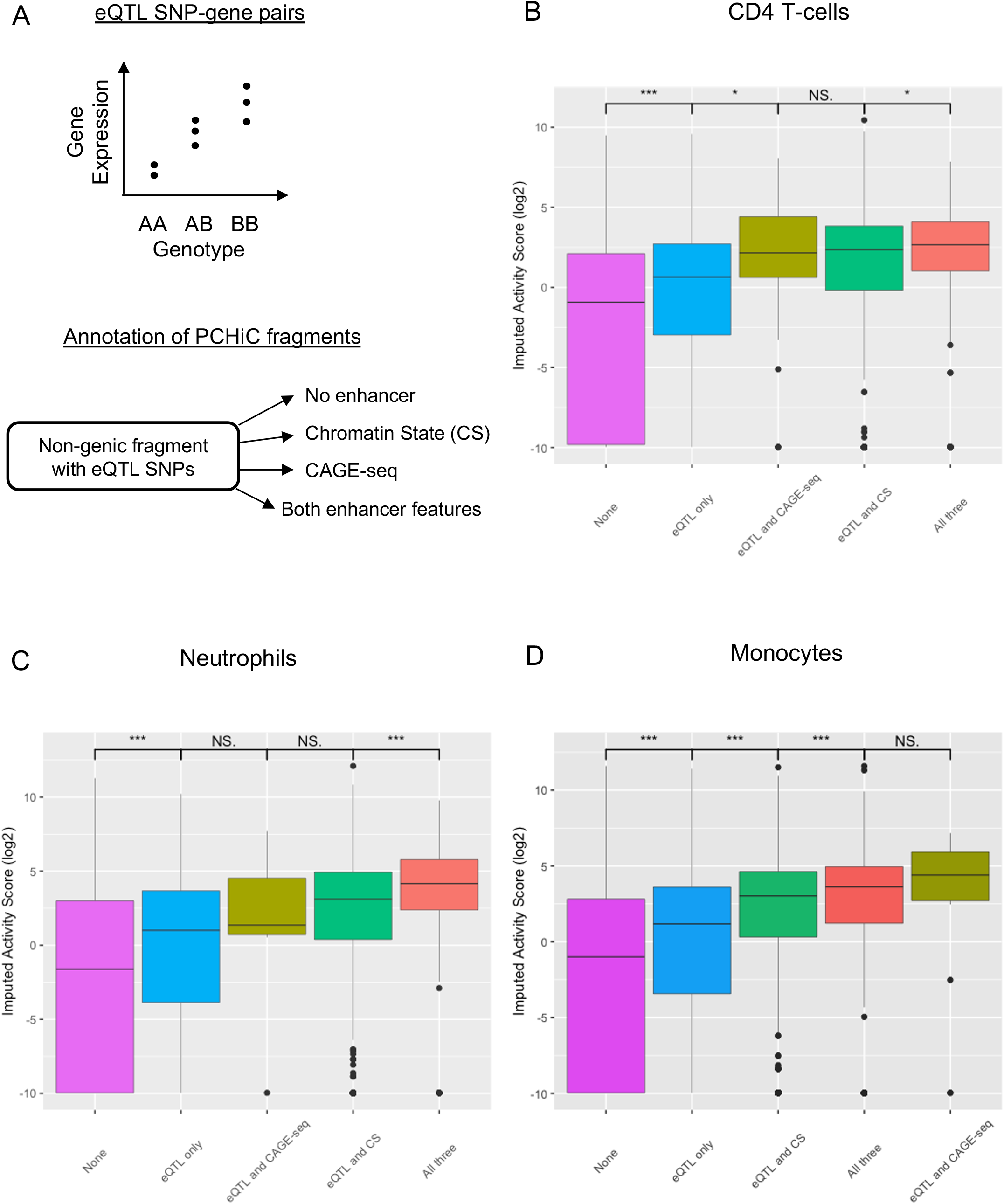
Enrichment of Esearch3D imputed activity score in non-genic regions with eQTL SNPs. (**A**) Schematic of how eQTL SNPs are detected and annotation of nodes in the human immune cells networks. Boxplots are ordered by the median of each group and asterisks indicate significance in Wilcoxon tests (* = p.value < 0.05, ** = p.value < 0.005 *** = p.value < 2.2e-16). (**B**) Imputed activity scores (IAS) in CD4+ T-cells. (**C**) Imputed activity scores in neutrophils. (**D**) Imputed activity scores in monocytes.

The BLUEPRINT Consortium generated eQTL data for three human immune cell types: monocytes, neutrophils and CD4+ T-Cells (20). We used this data to test the ability of Esearch3D to prioritise the regulatory potential of non-genic regions harbouring eQTL SNPs. For these cell types, we used available PCHi-C data (50) to generate 3D genome networks and annotated the non-genic nodes with the cell type specific eQTL SNPs, enhancer chromatin states from BLUEPRINT (51) and bi-directional CAGE-seq sites from the FANTOM project (16, 46). We found 46,078 monocyte nodes, 39,930 neutrophil nodes and 48,016 T-cell nodes with eQTL SNPs but only 41% of eQTLs in monocytes overlapped with nodes containing either FANTOM or BLUEPRINT enhancers with 28% in neutrophils and 31% in T-cells.

Interestingly, we observed a significantly higher IAS in non-genic nodes with eQTLs compared to non-genic nodes without eQTLs in all three cell types (p-value < 2.2e-16), even in nodes that did not overlap with experimentally defined enhancer features (**Figure 4B-D**). These eQTLs may point to loci that contain weaker enhancers that are not annotated by either CAGE-seq or chromatin state data. However, Esearch3D also predicts higher activity (IAS) when the non-genic nodes with eQTL SNPs also overlap with enhancer features (p-value < 2.2e-16) (**Figure 4B-D**). These results suggest that the IAS calculated by Esearch3D can help to prioritise eQTL SNPs with higher potential of being regulatory regions.

In line with previous results in mESC derived networks, we also find that IAS is generally higher in nodes with all three annotations. This further highlights the generalisability of our model to additional cell-types, including primary cell types that are difficult to assay for enhancer annotations that depend on transfections, such as CRISPR deletion/silencing experiments or STARR-seq screens.

### IAS can be used in conjunction with centrality measures to classify enhancer nodes

Using four commonly used network measures we calculated different topological characteristics of intergenic nodes in the mESC DHS-CHiC network (**Supplementary Table 3**). We calculated the degree, which reflects the number of edges, or connections, that the nodes in the network maintain. The betweenness centrality, which measures the number of shortest paths that pass through a node; this measurement reflects how important a node is in connecting other nodes. The closeness centrality that reflects how central a node is in the network relative to all other nodes. Finally, the eigenvector centrality, which measures the influence of a node within the network; this is calculated relative to the scores of other nodes in the network where a node with a high eigenvalue score connects to many other nodes with high eigenvalue scores.

A Random Forest classifier was then used to combine these scores along with IAS in order to classify intergenic enhancer nodes from the mESC DHS-CHiC network. We utilised the out-of-bag (OOB) error as a measurement to evaluate the contribution of each of the five topological metrics along with IAS when classifying intergenic enhancer and intergenic non-enhancer nodes. OOB measures the loss of classification performance of the random forest classifier when the feature of interest is removed. For capture Hi-C there is a bias between baits (such as promoters or, in this case, DHS sites) and other ends (non-bait fragments) due to the enrichment protocol (**Figure 5B**). This results in artificially lower connectivity for other end nodes when compared to baits (32). Due to the unique nature of capture Hi-C networks we evaluated the five topological metrics and IAS for non-genic baits and other ends separately, using two different Random Forest models.

**Figure 5.**
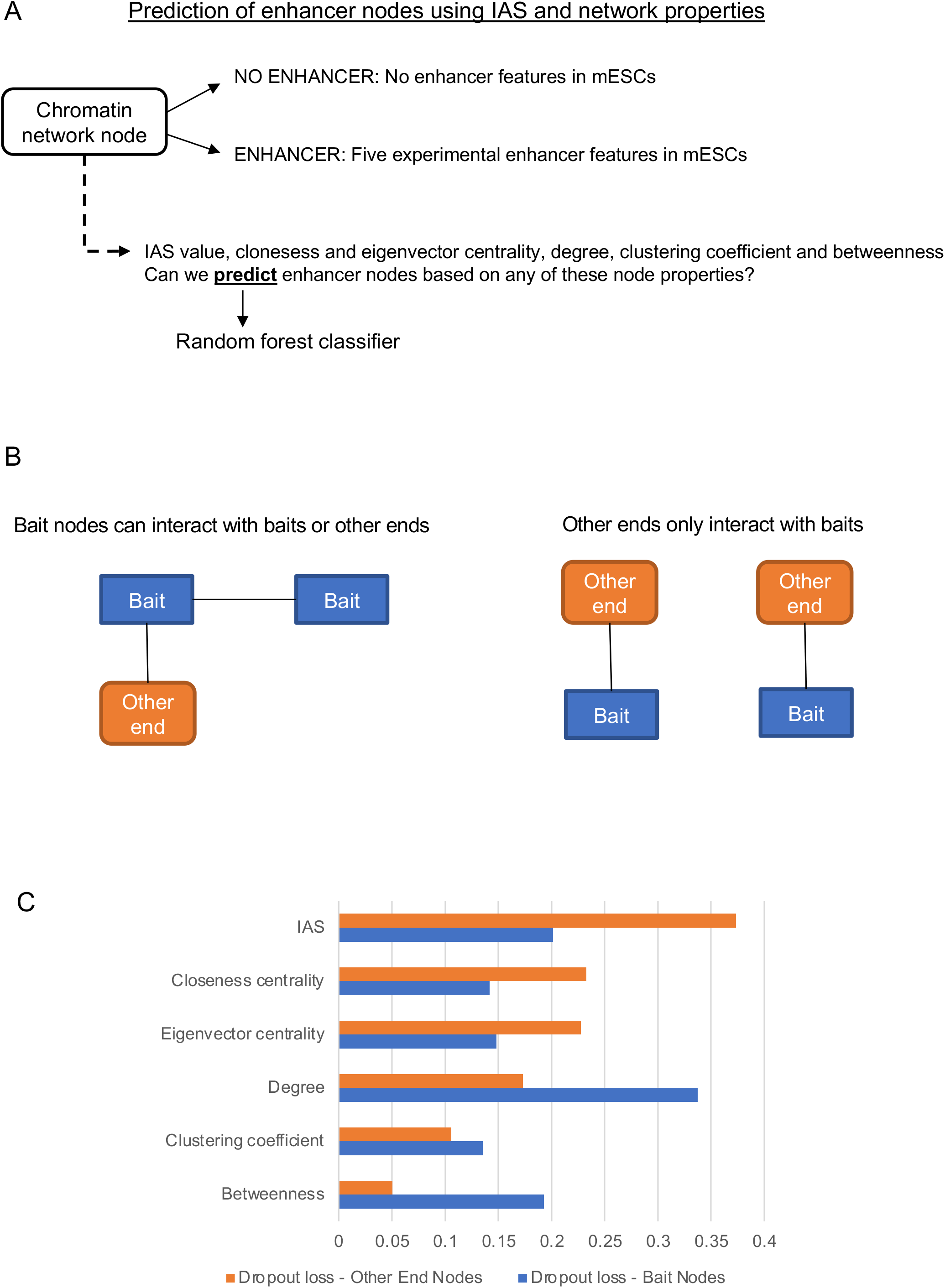
Esearch3D imputed activity score (IAS) and other network metrics can be used to predict enhancers. (**A**) Approach to classify enhancer and non-enhancer nodes. (**B**) Cartoon illustrating the different connectivity of bait (blue) and other end (orange) nodes in capture HiC, resulting in different network metrics. (**C**) The dropout loss of topological network measures used to build a Random Forest classifier of enhancer nodes (predicting the presence of at least one enhancer feature) with baits (blue) and other end (orange) nodes of the DHS-CHiC network.

We found that for non-genic baits, the degree centrality was the feature that was the most informative when classifying nodes into intergenic enhancers and intergenic non-enhancers with a drop out loss of 0.33 (**Figure 5C**). IAS was the next best measure with a dropout loss of 0.20. For other ends the best performing feature was IAS with a drop out loss of 0.37 (**Figure 5C**). Interestingly, the degree centrality was only the 4th most important feature for the other ends where connectivity information is lost due to the capture HiC design. In summary, IAS performs well for both types of fragments and contributes to enhancer prediction, highlighting the advantage of integrating gene expression data compared to only measuring the topological characteristics of intergenic nodes. This approach is particularly powerful to predict enhancers in less connected other end nodes, suggesting that the Esearch3D approach is a useful tool to compensate for the interaction sampling bias in capture HiC experiments.

### Esearch3D is implemented as an R package with shiny-based graphical user interface to explore the results

We have shown how IAS can be used to predict the location of enhancers. We have implemented Esearch3D as an R package. The method can be applied following the provided workflow with the chromatin interaction network and the gene expression profiles of the user’s cell-type of interest (**Supplementary Figure 3**). The software begins by performing the function “rwr_OVprop” twice. During the first step, the expression of the genes is propagated from their corresponding nodes into the genic nodes. In the second step, the gene expression is then propagated across the rest of the network. This imputes a proxy for transcriptional activity, IAS, at intergenic nodes.

The user can visualise the propagation results in a matrix format or with the graphical user interface (GUI) developed with R shiny (**Figure 6**). The GUI allows one to explore a gene expression profile after a network-based propagation, to investigate the imputed activity scores obtained by specific genes and their neighbourhoods and to import the propagated chromatin network into Cytoscape (52). The GUI allows users to visualise and focus on specific modules of the chromatin interaction network based on their interests. To illustrate the GUI functionality, we loaded the propagation results of the mESC DHS-CHiC network and explored the interactors of the gene node of the proto-oncogene *Myc* (**Figure 6**). The interaction (in orange) between the *Myc* node and node frag54325 indicates that the *Myc* locus is located inside this genomic fragment in mouse chromosome 15; the purple edges show the direct fragment-fragment interactions of the fragment containing *Myc*. The interacting fragment with the highest IAS value is frag54325 (chr15:61,807,659-61,821,282): this corresponds to a genomic region downstream of *Myc* that overlaps with all enhancer features (STARRseq, FANTOM bidirectional CAGE-seq, P300, enhancer chromatin states and RNAPII s2p. Interestingly, this genomic region overlaps with the Gsdmc gene cluster that is downstream of the recently discovered ‘blood enhancer cluster’, conserved between human and mouse, that controls *Myc* in mouse haematopoietic stem cells and progenitors (53). The region containing the GSDMC gene cluster has been shown to form part of the regulatory network of *MYC* in human cell lines (54), suggesting that this could be an important regulatory node in mESCs too.

**Figure 6.**
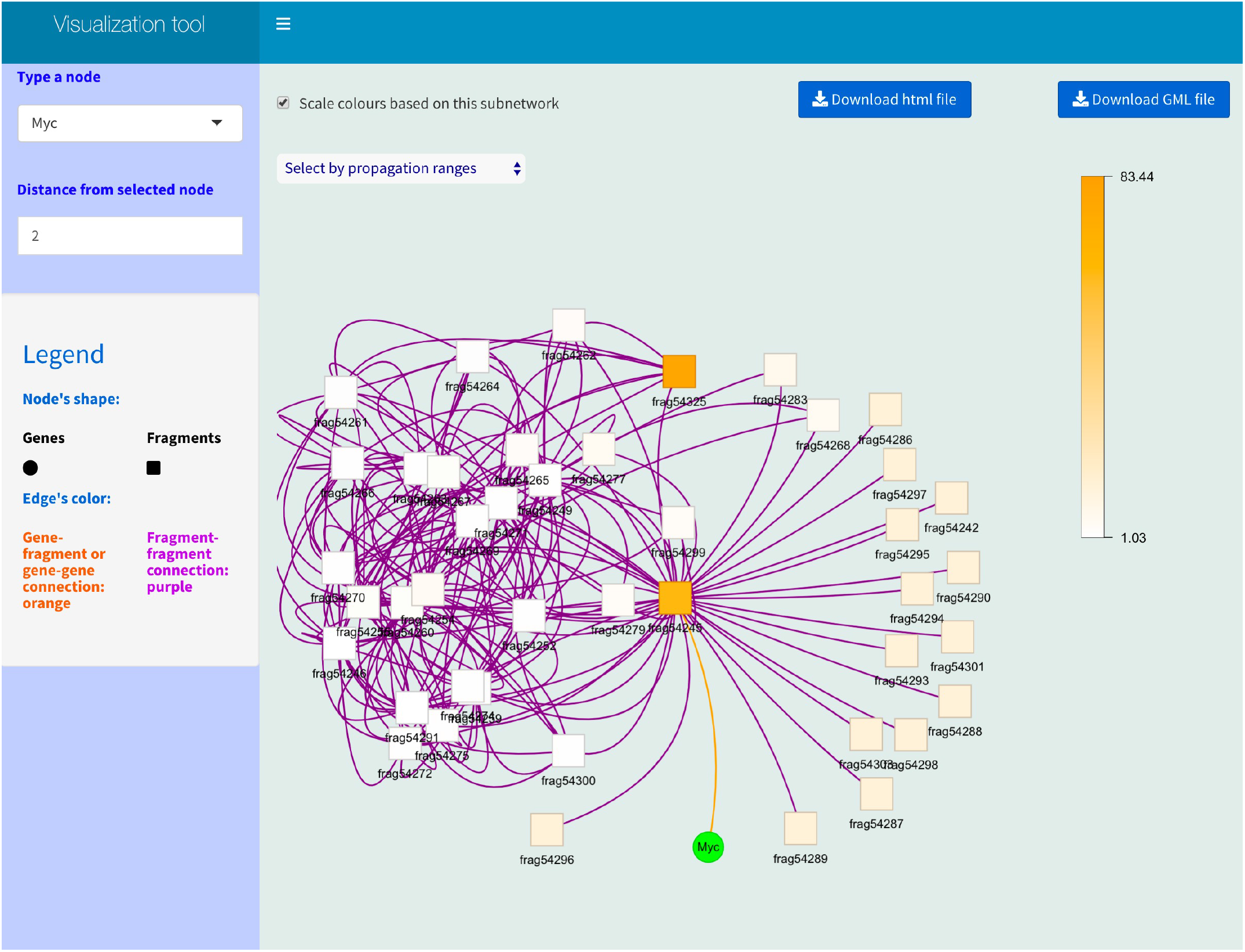
Visual exploration of Esearch3D results. The R shiny app provides a graphical user interface to explore sub-networks with the propagation results, which can be exported in Cytoscape format. The screenshot shows the direct interaction of *Myc* gene (represented as a node in the network, as explained in Fig. 2) with the DHS-PCHiC fragment containing it and direct interactors of that fragment (plus their interactions). The white to orange legend correspond to the Esearch3D IAS value of the fragments (scaled based on the visualised sub-network).

## Discussion

The folding of chromatin into hierarchical structures ultimately acts to localise genes with the prerequisite machinery for transcription such as RNAPII (32, 55) and the appropriate chromatin modifications (56). Whether gene expression dictates the chromatin architecture or vice-versa is poorly understood. In either case, it is clear that the organisation of chromatin plays an important role in mediating the communication between genes and enhancers. In this paper, we have outlined a method that can be used to predict enhancers by leveraging the relationship between gene expression and the global chromatin architecture. To achieve this we model interaction data from HiC-derived methods as networks.

We use a network propagation algorithm based on random walks to integrate gene expression data within the chromatin interaction network. The use of network propagation has been proposed previously in other fields, such as heat diffusion processes, energy states within electrical circuits, and graph kernels in machine learning (57). Other applications in biology include gene function prediction, module discovery, and drug target prediction, see (57). In cancer research for example, network propagation has been used to identify previously unknown proteins involved in the progression of cancers by propagating mutational frequencies across a protein-protein interaction network (58). We use the same concept with chromatin-chromatin interaction networks and propagating gene expression values instead of mutational frequencies. By doing so, Esearch3D is able to impute an activity score (IAS) in intergenic nodes by propagating the gene expression values from genic nodes across the chromatin interaction network. IAS is a metric that summarises the importance of an intergenic node in the context of its proximity to all of the genic nodes within the network, the relative expression of those genes as well as the topology of the entire network. Most widely used methods to identify enhancers do not incorporate gene expression, nor do they evaluate the subtle, but important influences exerted by chromatin’s global organisation. Esearch3D builds on this modelling 3D genome data as networks and then exploits graph-theory algorithms to integrate RNA-seq data in order to calculate an imputed activity score (IAS).

The underlying assumption of Esearch3D is that the activity, defined by the gene expression, of one chromatin fragment is influenced by, or can influence, the activity of all other chromatin fragments. Transcriptional factories occur through dynamical processes that lead to phase separation (47). These are, in part, influenced by relative levels of gene expression across the chromatin interaction network suggesting a level of global influence at a local level (59). We too identify such behaviour in our model. We propagate the expression of all of the genes simultaneously in order to model the global interdependent relationships between genes, chromatin conformation and enhancers. Indeed, we find that the performance of Esearch3D is driven by propagating the expression of all genes, whereas a poor performance is seen when propagating the expression of a single gene. This would suggest that gene expression and chromatin organisation are linked at a global level.

The challenges that exist in finding enhancers stem from a fundamental gap in our knowledge about their composition and mode of action. It is becoming increasingly apparent that chromatin architecture plays an important role in gene regulation by localising enhancers to gene promoters in specific patterns. This idea is commensurate with the looping mechanisms that localise enhancers and gene promoters resulting in an increased contact frequency (3). These effects can be obvious as shown by the loss of interaction between the ZRS enhancer and the *SHH* promoter following the deletion of a key CTCF site (60). This suggests that enhancers and genes are distinctly connected within the 3D chromatin structure compared to other loci. However, to what extent 3D interactions are always essential for gene regulation is debatable. Some studies have shown that the disruption of some TADs does not lead to wide scale changes to gene expression (61) suggesting that mechanisms other than the chromatin topology may regulate gene expression. Alternatively, complex networks are known to be robust to attack (62) and, as such, deletion of interactions within the network should not always be expected to result in large gene expression changes. Other explanations such as redundancy in the enhancer networks are equally plausible (6, 7). In such cases, more care should be taken when making claims about the role of chromatin organisation in gene regulation when regions are studied in isolation. In fact, we show that by studying the relationships between gene expression and the global structure of chromatin we can accurately identify enhancer regions. We have shown that enhancers and genes within our networks tend to contain distinct topological features as shown by the centrality scores when compared to non-enhancer and intergenic nodes and that these features can be used to predict intergenic enhancer nodes. Our data would therefore suggest that chromatin organisation plays an important role in coordinating gene regulation.

## Materials and Methods

### From 3C to Networks

Processed 3C datasets were downloaded for mouse embryonic stem cells (mESCs, serum) and three immune human cell types. For mESCs, we used DNaseI capture Hi-C dataset from Joshi et al (42) and promoter-capture HiC from (63) and reprocessed by Pancaldi et al., (32), both aligned to the GRCm38/mm9 reference genome. For the human primary immune cells, we used data from Javierre et al., aligned to the GRCh37/hg19 reference genome (50). Contacts were normalised and loops were called in the three datasets using the CHiCAGO algorithm (64). The normalised contact matrix was then transformed into an unweighted and undirected network *G* = (*V,E*) where the chromatin fragments are represented as nodes *V* and their interactions as edges *E*.

### Gene annotation

Each node in the network represents a distinct genomic locus each containing biological features that infer their functional properties. To identify intergenic enhancers each node was annotated with genes from Ensembl version 75 (65).

### Enhancer annotation

As is common in all enhancer prediction models, the identification of a suitable ‘truth set’ of validated enhancers is non-trivial. Currently gold standard approaches such as CRISPR perturbation are limited in both the scalability to assess putative enhancer sequences genome wide as well as the availability of this data for multiple cell-types. As there is no genome-wide gold standard for enhancer identification in mESCs, we identified several enhancer associated features that could be used to label the intergenic nodes as enhancers and non-enhancers.

We first used genome wide chromatin profiles to identify putative enhancers in mESCs based on histone marks, including H3K27ac, H3K4me1 and H3K4me3 (**Supplementary Table 4**). To efficiently summarise the data we utilised the popular ChromHMM method (66) and only bins with a posterior probability above 0.95 were considered (44). The nodes were then labelled with the chromatin state bins and the percentage overlap of each state was calculated per fragment. This was further supplemented with the annotation of enhancer associated features from experimental data; these included three RNAPII variants and P300 ChIPseq (44), STARR-seq (45) and bi-directional CAGE-seq sites from the FANTOM project (16, 46). By combining these data we were then able to distinguish both intergenic nodes as enhancers as well as stratify this set into high and low confidence groups based on the number of features present. For the immune cells, we defined putative enhancers using chromatin states (**Supplementary Table 5**) from BLUEPRINT (51), eQTL SNPs (20) and bi-directional CAGE-seq sites from the FANTOM project (16, 46).

### Gene expression data

Normalised gene expression single-cell RNAseq data for mESCs was downloaded from ESpresso (67) (https://espresso.teichlab.sanger.ac.uk), where we took the mean expression per gene of the cells in serum. The expression of CD4 T-cells, monocytes and neutrophils was obtained from BLUEPRINT population dataset (20). The un-normalised read counts where re-normalised separately for each cell type using DESeq2 (68), keeping the chromosome X (in the original publication, X and Y chromosomes were removed before normalisation and the three cell types were normalised together). Finally, we took the mean gene expression all the individuals.

### Development of a network based expression propagation strategy

Our network-based propagation algorithm, implemented within Esearch3D, uses the propagation function *f*-defined by Zhou et al. (41). This algorithm is best understood as simulating a process where node A in the network contains an *a priori* value. In the first iteration, the propagation algorithm spreads or ‘propagates’ this value to the adjacent neighbours of A; which are B, C and D. In the second iteration B, C and D can propagate their values from the first iteration to their adjacent neighbours E and F (**Supplementary Figure 1**). This process continues for a set number of iterations based on the user input. These processes have a wide range of applications, but fundamentally they are all able to impute information in nodes with no *a priori* value based on the starting value at A. Another feature of this algorithm is that the propagation processes factors in the global topology of the network. The number of connections between the nodes and their patterns will ultimately influence the final imputed value of each node. This can be useful when we want to assess the importance of the topological characteristics of nodes in the chromatin interaction network in relation to gene expression. In our case, we wanted to understand if enhancers were uniquely connected within the network to transmit regulatory information to genes. To test this we calculated an **imputed activity score (IAS)** in intergenic nodes based on the topology of a chromatin interaction network using gene expression as our *a priori* starting values in gene nodes.

### Network-based propagation algorithm

Given a connected weighted graph G(V, E) with a set of nodes V={v1, v2,…, vN} and a set of links E={(vi, vj)|vi, vj∈V}, a set of source/seed nodes S⊆V and a N×N adjacency matrix W of link weights equal to 1 (**Supplementary Figure 1**). Here, we use a Random Walk with Restart (RWR) algorithm for measuring relative importance of node vi to S. RWR mimics a walker that moves from a current node to a randomly selected adjacent node or goes back to source nodes with a back-probability γ∈(0, 1). RWR can be formally described as follows:

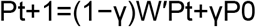

where Pt is an N×1 probability vector of |V| nodes at a time step t of which the ith element represents the probability of the walker being at node vi∈V, and P0 is the N×1 initial probability vector W’ is the transition matrix of the graph, the (i,j) element in W’, denotes a probability with which a walker at vi moves to vj among V{vi}.

### Esearch3D R package

Esearch3D is implemented in R, available at https://github.com/InfOmics/Esearch3D. It takes as input a chromatin interaction network and a cell-type specific gene expression profile. The network must be divided in two components, one composed of genes and the 3C fragments they map to where the overlap is represented as an interaction. The second includes the interactions (contacts) from the 3C data. The network can be inputted as an adjacent matrix, edge list or *igraph* object (https://igraph.org). The gene expression profile must be in the standard matrix format Nx1 such that rows are genes and columns are the replicates, conditions. Esearch3D propagates the gene expression values into genic and not genic fragments of the network in parallel using the R package *doParallel* (version 1.0.17). The functions “create_net2plot” and “start_GUI” represent results as a R *igraph* object and visualise it in a *shiny* (version 1.7.2) GUI.

Esearch3D enables researchers to investigate genes or fragments by providing the names of the genes to analyse and running a new propagation with the function called “rwr_SGprop”. This step provides how much each gene of interest contributed to give information to the chromatin network nodes (genomic fragments) and also how much each fragment received information from the genes of interest. Esearch3D documentation includes a vignette detailing the workflow to classify any node as an enhancer or not, where information about the number of enhancer annotations associated with the fragments of the chromatin interaction network are made available. This workflow enables the user to build a machine learning algorithm to classify nodes within the network as enhancers. Esearch3D computes the centrality measures of the nodes with the function “get_centr_info”, runs the multi-gene propagation and combines all the information in a matrix format. The subsequent matrix contains, for each node in the network, the IAS, the provided centrality measures as well as the number of enhancer annotations (provided by the user). The user can then build the classifier with the function “enhancer_classifier”. The function “train”, tunes and tests a random forest classifier with the R package mlr3. The best classifier can then be passed to the function “explain_classifier” that builds an explainer with the R package DALEX. The explainer describes what the classifier learnt, how much each feature contributes to the prediction and which decisions are used to predict an unlabelled node. Topological features are calculated with *igraph* R package and the Random Forest models are built with *DALEX* R package (version 2.4.1) - see vignettes at https://github.com/InfOmics/Esearch3D for details.

### Precision-recall curves

Precision-recall curves plot, at increasing thresholds, the precision, which is the fraction of nodes correctly predicted as enhancers (i.e. positive for the five enhancer features in **Figure 3A**), and the recall, which is the fraction of the total number of enhancers predicted. Often, a tradeoff occurs between these two metrics whereby increased recall reduces the precision and vice versa. When plotted on the same graph the area under the curve produced by this plot indicates the general classification performance of the model. The precision and recall of enhancer nodes using centrality measures and the corresponding area under the curve (AUPRC) were calculated using the *precrec* library (version 0.12.7) (69). The baseline values were calculated as the total number of enhancer nodes as a proportion of the total number of nodes in the chromatin interaction network.

## Acknowledgements

We thank Caelinn James and Wilhelmina Lucinescu for testing the initial versions of ESearch3D, and Lu Chen, Nicole Soranzo, Simone Ecker and Immaculada Hernandez-Lopez for their help re-processing the BLUEPRINT RNAseq data of immune cells. We also thank Vera Pancaldi and Biola Javierre for their useful comments that helped us to improve the text.

This work was supported by the Wellcome Trust (206103/Z/17/Z to D.R. and Newcastle University ISSF to fund the living expenses of L.G. during his internship). M.H. was supported by an EPSRC DTP (Biological Informatics) studentship.

