## Supplementary material for "Esearch3D: Propagating gene expression in chromatin networks to illuminate active enhancers": Suplementary Figures and Tables

A

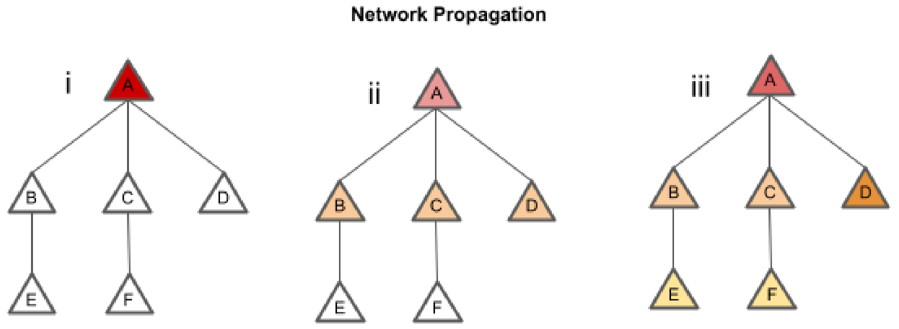


B

**Adjacency Matrix**

| A | B | C | D | E | F |
|---|---|---|---|---|---|
| 0 | 1 | 1 | 1 | 0 | 0 |
| 1 | 0 | 0 | 0 | 1 | 0 |
| 1 | 0 | 0 | 0 | 0 | 1 |
| 1 | 0 | 0 | 0 | 0 | 0 |
| 0 | 1 | 0 | 0 | 0 | 0 |
| 0 | 0 | 1 | 0 | 0 | 0 |

**Transition Matrix**

| A | B | C | D | E | F |
|---|---|---|---|---|---|
| 0 | 0.5 | 0.5 | 1 | 0 | 0 |
| 0.3 | 0 | 0 | 0 | 1 | 0 |
| 0.3 | 0 | 0 | 0 | 0 | 1 |
| 0.3 | 0 | 0 | 0 | 0 | 0 |
| 0 | 0.5 | 0 | 0 | 0 | 0 |
| 0 | 0 | 0.5 | 0 | 0 | 0 |

**Expression**

| | | | | | | |
|---|---|---|---|---|---|---|
| 100 | 0 | 0 | 0 | 0 | 0 | $t_0$ |
| 94 | 6 | 6 | 6 | 0 | 0 | $t_1$ |
| 95.8 | 4.8 | 4.8 | 5.4 | 0.6 | 0.6 | $t_2$ |

**Supplementary Figure 1.** A) i) A toy model of network propagation run with two iterations with an insulating score of 0.2. A single gene node A contains an a priori expression value. aii) In the first iteration, the value from A is smoothed to its direct neighbours. aiii) In the second iteration the values are further smoothed and important nodes such as B and C are ranked higher. B) i)This process is achieved by representing the interactions between nodes of the network in an adjacency matrix. ii) The degree of each node is represented in the diagonal matrix. iii) The probabilities of transitioning from one node to another is calculated from the diagonal matrix and the adjacency matrix using equation 1. These results are then represented in the degree-normalised transition matrix. iv) The expression value for each node at the start and after each of the two iteration values after propagation using equation 2.

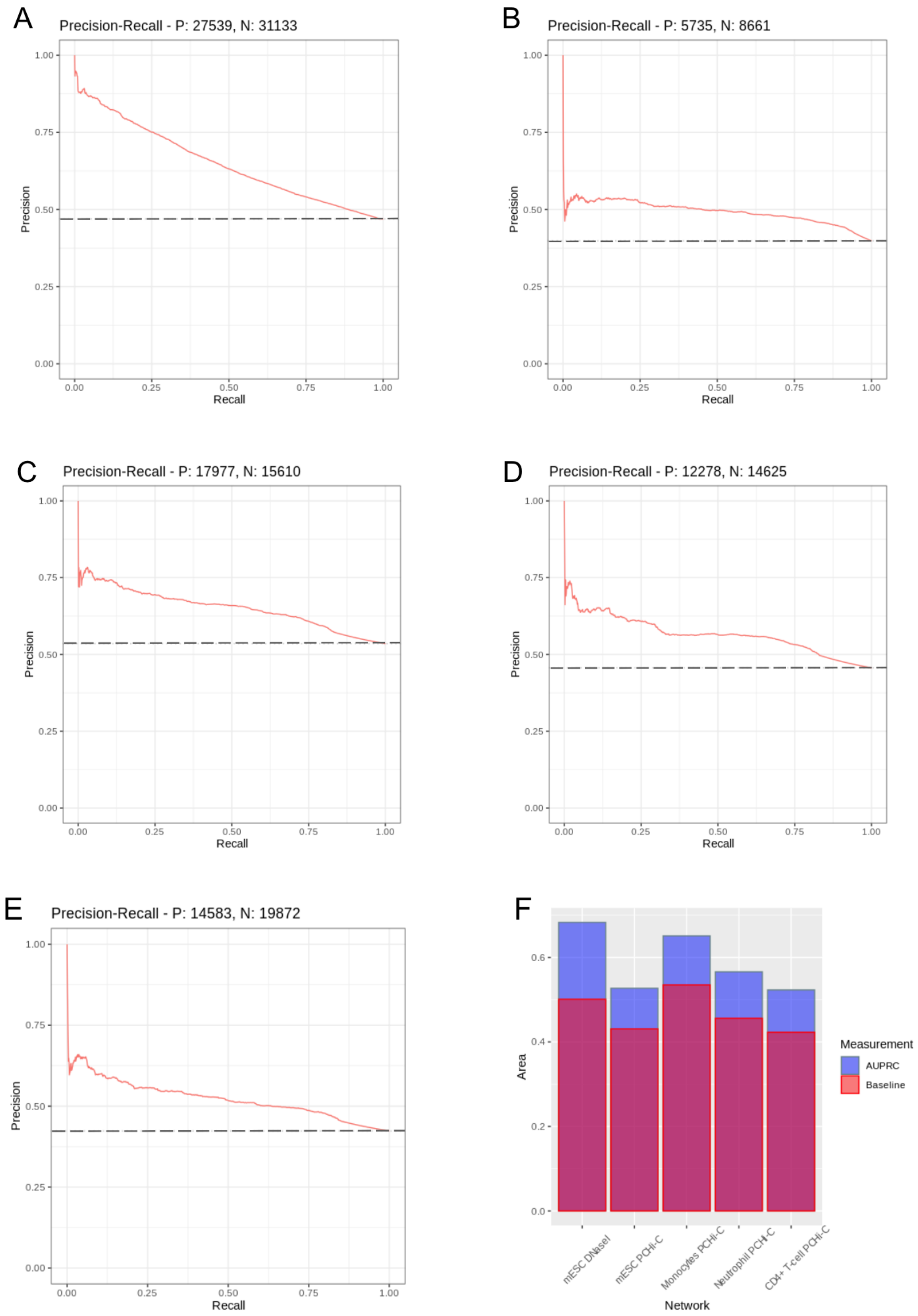


**Supplementary Figure 2. Precision recall curves measuring the overall performance of IAS in the prediction of enhancer nodes containing a minimum of 1 enhancer feature.**

(A) mESC DNaseI derived network AUPRC = 0.650 over a baseline of 0.469. (B) mESC PCHi-C derived network AUPRC = 0.493 over a baseline of 0.398. (C) Monocytesderived network AUPRC = 0.0.651 over a baseline of 0.535. (D) Neutrophil derived network AUPRC = 0.566 over a baseline of 0.456. (E) CD4+ T-Cell derived network AUPRC = 0.0.523 over a baseline of 0.423. (F) An overlay barchart showing the AUPRC (blue) relative to the baseline score (red) for each of the the precision recall curves.

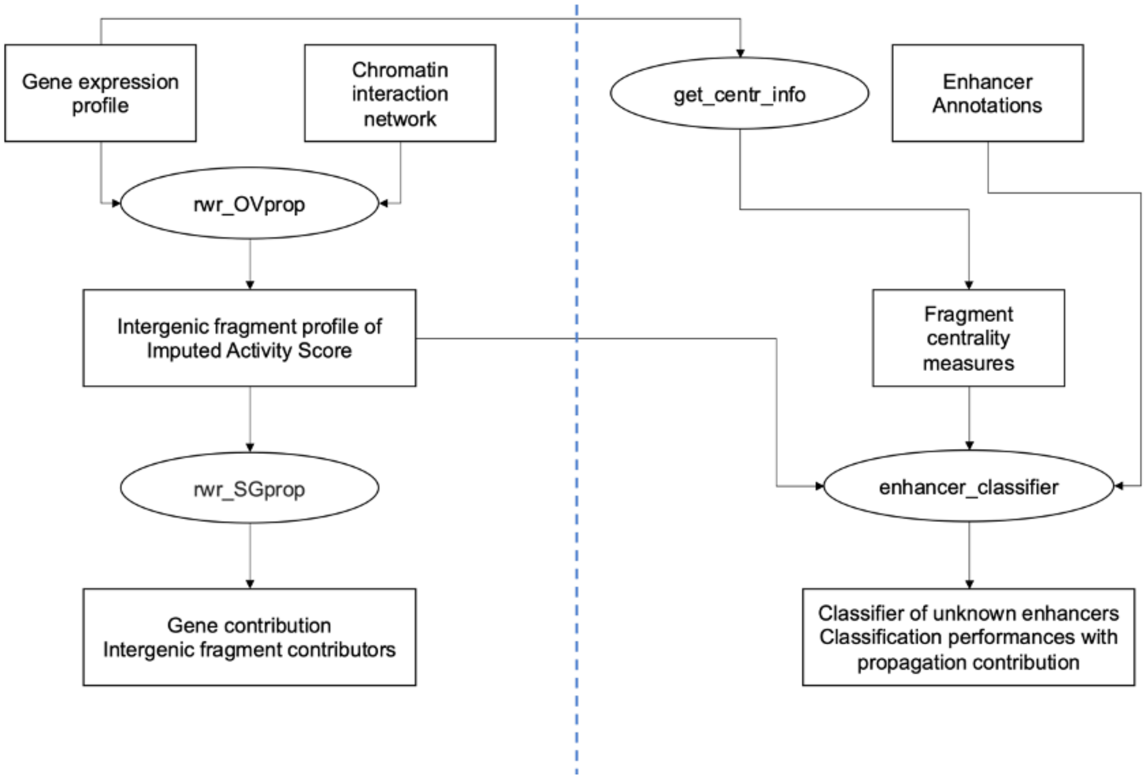


**Supplementary Figure 3. Pipeline implemented in the Esearch3D R package.** Package and documentation available at https://github.com/InfOmics/Esearch3D.

| | Intrachromosomal Interactions | Interchromosomal interactions | Connected components | % nodes in the LCC | Density |
|---|---|---|---|---|---|
| Monocytes (PCHi-C) | 170,329 | 1,101 | 1938 | 34.71% | 0.000037 |
| Neutrophils (PCHi-C) | 146,146 | 1,049 | 2136 | 32.26% | 0.000045 |
| CD4+ T-Cells (PCHi-C) | 218,377 | 1,062 | 1375 | 51.63% | 0.000038 |
| mESC (DNaseI) | 782,466 | 9,457 | 776 | 98.12% | 0.000060 |
| mESC (PCHi-C) | 69,847 | 2,374 | 4073 | 66.23% | 0.000048 |

**Supplementary Table 1.** Summary statistics (part 1) of the five capture HiC derived networks used. LCC: Largest connected component.

| | Total number of nodes | Total number of edges | Average degree |
|---|---|---|---|
| Monocytes (PCHi-C) | 96,442 | 171,430 | 3.56 |
| Neutrophils (PCHi-C) | 81,242 | 147,195 | 3.62 |
| CD4+ T-Cells (PCHi-C) | 106,848 | 219,439 | 4.11 |
| mESC (DNaseI) | 162,615 | 791,903 | 9.69 |
| mESC (PCHi-C) | 53,920 | 72,221 | 2.61 |

**Supplementary Table 2***.* Summary statistics (part 2) of the five capture HiC derived networks used .

| Enhancer (Non-Enhancer) | Degree | Harmonic Closeness | Betweenness | Eigenvector |
|---|---|---|---|---|
| Monocytes (PCHi-C) | 2.36 (1.86) | 222 (207) | 3526 (3443) | 0.000757 (0.000175) |
| Baits | 13.61 (8.90) | 195 (174) | 43,421 (12,380) | 0.001378 (0.000000) |
| OE | 1.89 (1.61) | 223 (207) | 1,864 (3,123) | 0.000731 (0.000182) |
| Neutrophils (PCHi-C) | 2.48 (1.95) | 191 (169) | 2662 (1318) | 0.000801 (0.000138) |
| Baits | 12.99 (8.45) | 166 (108) | 22,997 (13,675) | 0.001591 (0.000487) |
| OE | 1.97 (1.66) | 193 (172) | 1,672 (771) | 0.000762 (0.000122) |
| CD4+ T-Cells (PCHi-C) | 2.71 (2.08) | 505 (474) | 7547 (4038) | 0.000768 (0.000131) |
| Baits | 14.76 (11.66) | 470 (389) | 70,396 (26,903) | 0.000404 (0.000523) |
| OE | 2.18 (1.80) | 506 (477) | 4,738 (3,372) | 0.000785 (0.000119) |
| mESC (DNaseI) | 10.62 (2.63) | NA | 571,553 (74,722) | 0.000161 (0.000053) |
| Baits | 17.48 (7.43) | NA | 983,475 (212,061) | 0.000203 (0.000080) |
| OE | 2.57 (1.99) | NA | 87,620 (56,720) | 0.000113 (0.000049) |
| mESC (PCHi-C) | 1.44 (1.20) | 913 (707) | 83,562 (33,001) | 0.000638 (0.000185) |
| Baits | 4.58 (2.53) | 619 (316) | 1,028,513 (292,101) | 0.000091 (0.000005) |
| OE | 1.43 (1.19) | 914 (709) | 78,829 (31,154) | 0.000641 (0.000186) |

**Supplementary Table 3.** Average centrality scores between enhancer and non-enhancer nodes (median between parentheses). OE: Other end.

| State | Associated Function | Histone Marks |
|---|---|---|
| State 1 (E1) | Transcription | H3K4me1 & H3K79me3 & H3K36me3 |
| State 2 (E2) | Transcription | H3K4me1 & 5hmC & H3K9ac & H3K4me2/3 H3K36me3 |
| State 3 (E3) | Transcription | H3K4me1 & 5hmC & H3K36me3 |
| State 4 (E4) | Transcription | H3K36me3 |
| State 5 (E5) | Transcription | H3K36me3 |
| State 6 (E6) | Heterochromatin | H3K9me3 & H4K20me3 |
| State 7 (E7) | Low Signal | N/A |
| State 8 (E8) | Heterochromatin | 5fC |
| State 9 (E9) | Low Signal | N/A |
| State 10 (E10) | Heterochromatin | 5hmC & 5mC |
| State 11 (E11) | Enhancer | H3K4me1 & 5hmC |
| State 12 (E12) | Enhancer | H3K4me1 &5hmC &H3K4me2/3 |
| State 13 (E13) | Enhancer | H3K4me1 |
| State 14 (E14) | Enhancer | H3K4me1 & H3K27ac & H3K9ac & H3K4me2/3 |
| State 15 (E15) | Active Promoter | H3K4me1/2/3 & H3K27ac & H3K9ac & H3K79me2 & H3K36me3 |
| State 16 (E16) | Active Promoter | H3K4me2/3 & H3K27ac & H3K9ac & H3K79me2 |
| State 17 (E17) | Active Promoter | H3K4me2/3 |
| State 18 (E18) | Bivalent Promoter | H3K4me1/2/3 & H3K9ac &5hmC & H3K27me3 |
| State 19 (E19) | Repressed Promoter | H3K27me3 |
| State 20 (E20) | Insulator | CTCF |

**Supplementary Table 4.** Summary table of chromatin states for mouse embryonic stem cells from Juan *et al* (44).

| State | Associated Function | Histone Marks |
|---|---|---|
| State 1 (E1) | Weak Transcription | H3K36me3 |
| State 2 (E2) | Transcription | H3K36me3 |
| State 3 (E3) | Heterochromatin | H3K9me3 |
| State 4 (E4) | Low signal | N/A |
| State 5 (E5) | Heterochromatin | H3K27me3 & H3K9me3 |
| State 6 (E6) | Heterochromatin | H3K27me3 |
| State 7 (E7) | Repressed Promoter | H3K4me1 & H3K4me3 & H3K27me3 |
| State 8 (E8) | Enhancer | H3K4me1 |
| State 9 (E9) | Enhancer | H3K4me1 & H3K27ac |
| State 10 (E10) | E-Promoter | H3K4me1 & H3K27ac & H3K4me3 |
| State 11 (E11) | Promoter | H3K27ac & H3K4me3 |

**Supplementary Table 5.** Summary table of chromatin states for human immune cells (monocytes, neutrophils and CD4 T-cells) from BLUEPRINT Consortium as described in Carrillo-de-Santa-Pau *et al* (51).
